# Unraveling the Potential of Hypericin: A Novel Approach to Targeting PCSK9 Mutations in Cholesterol Management

**DOI:** 10.1101/2024.10.06.616901

**Authors:** Ivan Vito Ferrari

**Affiliations:** University of Rome Tor Vergata, Department of Industrial Engineering, Via del Politecnico 1, 00133 Rome, Italy

**Keywords:** PCSK9, LDLR, hypericin, molecular docking, in silico mutational analysis, cholesterol metabolism, natural compounds, mutations, hypercholesterolemia

## Abstract

Proprotein convertase subtilisin/kexin type 9 (PCSK9) plays a pivotal role in regulating cholesterol metabolism by promoting the degradation of low-density lipoprotein receptors (LDLR), leading to elevated levels of LDL cholesterol in the bloodstream. Recent studies have identified key residues—Arg-194, Asp-374, and Phe-379—that are crucial for the interaction between PCSK9 and LDLR, and mutations in these residues can significantly alter this interaction. In this study, we employed molecular docking and in silico mutational analysis to assess the binding affinities of natural compounds, including Hypericin, amentoflavone, Bilobetin, Gingetin, and Progesterone, to wild-type and mutant PCSK9. We explored nine specific PCSK9 mutations (R194, D374Y, F379), targeting changes in residues known to influence LDLR binding affinity. Our results indicate that Hypericin demonstrates favorable binding to wild-type PCSK9, interacting with the critical residues involved in LDLR binding, while other mutations yield diverse effects on compound affinities. Notably, certain mutations, such as R194A and D374Y, significantly impact binding affinities, revealing both potential benefits and challenges in targeting PCSK9 with natural compounds. This study provides insights into potential therapeutic strategies for modulating PCSK9 activity and suggests new avenues for developing safer, more effective treatments for hypercholesterolemia.

## Introduction

Hypercholesterolemia refers to elevated cholesterol levels in the blood, which poses a significant risk to health due to its role in the development of atherosclerosis, increasing the likelihood of cardiovascular diseases such as heart attack and stroke. Cholesterol is vital for cell membrane formation and various metabolic processes, including the synthesis of vitamin D, bile salts, and hormones like testosterone and cortisol. However, excessive cholesterol, particularly low-density lipoprotein (LDL), often called “bad cholesterol,” can accumulate in the arterial walls, leading to atherosclerosis[1–5]. On the other hand, high-density lipoprotein (HDL) helps remove cholesterol from the bloodstream, transporting it to the liver for processing and elimination, hence earning the reputation of “good cholesterol.” Elevated LDL levels are strongly associated with cardiovascular events, while high HDL levels are linked to reduced cardiovascular risk. Familial hypercholesterolemia and combined hyperlipidemia are genetic conditions that cause abnormally high cholesterol levels, often leading to early cardiovascular disease. In contrast, secondary causes, such as a sedentary lifestyle, high-fat diets, diabetes, and other medical conditions, are more common contributors to hypercholesterolemia. Understanding these mechanisms is essential in managing cholesterol levels to mitigate the risk of cardiovascular diseases [1–5].

Hypercholesterolemia is a significant risk factor for cardiovascular diseases, and PCSK9 (Proprotein convertase subtilisin/kexin type 9) plays a critical role in regulating LDL-C levels through the degradation of LDLR. Mutations in the PCSK9 gene, such as Asp-374-Tyr, increase the affinity of PCSK9 for LDLR, contributing to elevated plasma LDL-C levels[6–9].

This study aims to explore the interactions between PCSK9 and potentially inhibitory compounds, analyzing binding affinity and interactions in the presence of specific mutations.

Proprotein convertase subtilisin/kexin type 9 (PCSK9) is a key regulator of low-density lipoprotein (LDL) metabolism in the body, primarily by modulating the degradation of hepatic low-density lipoprotein receptors (LDLRs). The interaction between PCSK9 and LDLR has significant implications for cholesterol homeostasis and cardiovascular health. Elevated plasma levels of LDL-C are a major risk factor for atherosclerosis and other cardiovascular diseases, making the understanding of the molecular mechanisms governing LDLR regulation essential[6].

PCSK9 binds to the epidermal growth factor-like repeat A (EGF-A) domain of the LDLR, leading to its internalization and subsequent degradation in lysosomes. This process results in a decrease in the number of LDLRs available on the cell surface, impairing the liver’s ability to clear LDL-C from the bloodstream. Genetic studies have identified various mutations in the PCSK9 gene that alter its function, with gain-of-function mutations, such as Asp-374-Tyr, leading to increased LDL-C levels and a heightened risk of cardiovascular diseases. Conversely, loss-of-function mutations have been associated with significantly reduced plasma LDL-C levels and a lower incidence of heart disease.

Given the critical role of PCSK9 in cholesterol regulation, it has emerged as a prominent target for therapeutic intervention. Monoclonal antibodies that inhibit PCSK9 have shown promise in clinical trials, significantly reducing LDL-C levels and providing a new avenue for patients who are statin-resistant or require additional LDL-C lowering. However, the development of small molecule inhibitors could offer advantages in terms of accessibility and administration, and an understanding of the interactions between PCSK9 and potential inhibitors is essential for drug design [6–9].

This study delves into the mechanisms behind cholesterol regulation, focusing on the role of PCSK9 and its interaction with LDL receptors (LDLR). Through in silico analysis, we investigate the effects of various natural compounds on PCSK9 mutations, offering potential insights for therapeutic interventions in hypercholesterolemia management.

In this study, we performed an in silico analysis [10,11] of the interactions between PCSK9 and several potential inhibitory compounds, including hypericin, amentoflavone, gingetin, bilobetin, and progesterone. Our focus was on examining how specific mutations in PCSK9 affect binding affinity and interaction profiles with these compounds. By analyzing the binding energy and molecular interactions across different mutations, we aim to provide insights into how these alterations might influence the therapeutic potential of various compounds in modulating PCSK9 activity.

## Materials and Methods

We performed a comprehensive virtual screening and molecular docking study to evaluate the binding affinity of 100 selected natural compounds against various PCSK9 variants. The workflow involved several key steps:

1. Protein Preparation: The structure of PCSK9 was initially obtained from the Protein Data Bank (PDB)[6]. To ensure accuracy in our analysis, the protein structure was repaired using the Protein Repair and Analysis Server (Osita Sunday Nnyigide et al., 2022) [12]. This server allows the addition of missing heavy atoms and hydrogen atoms, alongside assigning secondary structures based on amide interactions. Once the protein was fully prepared, we minimized its structure using the SPDBV (Swiss-Pdb Viewer) to optimize its geometry for docking [13].
2. Docking Procedure: Docking simulations were conducted using the PyRx software suite (version 0.8), known for its utility in molecular docking and virtual screening applications. The docking analysis focused on the key residues of PCSK9—Arg-194 (R194), Asp-374 (D374), and Phe-379 (F379) based on the critical findings of Kwon et al. (2008) [6]. These residues are essential for PCSK9’s interaction with the epidermal growth factor-like repeat A (EGF-A) domain of the LDLR, facilitating its degradation. To evaluate the binding affinities, we screened 100 natural compounds, calculating their binding energies (expressed in kcal/mol) to determine potential inhibitors.
3. Mutation Analysis: After performing docking on the wild-type PCSK9, we mutated the three main residues of interest (R194, D374, and F379) using Chimera’s Rotamer Function (Shandler, 2010) [14]. This tool allowed us to introduce mutations systematically in the PCSK9 structure, generating different variants to assess the impact on binding affinity and interactions with the compounds. These mutations were designed to mimic gain-of-function and loss-of-function scenarios commonly found in PCSK9 studies.
4. Visualization and Interaction Analysis: Following the docking experiments, we visualized the protein-ligand interactions using PyMOL [15] and the Discovery Studio Biovia Visualizer [16]. These tools enabled us to analyze the binding poses and determine key molecular interactions, including hydrogen bonds, hydrophobic interactions, and other non-covalent forces that stabilized the compounds within the active site of PCSK9. Through this in silico approach, we systematically assessed how natural compounds interacted with both the wild-type and mutant forms of PCSK9, providing insights into their potential as inhibitors to reduce LDL-C levels in hypercholesterolemic conditions.

The mutations were analyzed in the following order:

· Wild Type (R194, D374, F379)

· Mutated 1 (R194, D374Y, F379)

· Mutated 2 (R194A, D374Y, F379W)

· Mutated 3 (R194, D374Y, F379W)

· Mutated 4 (R194, D374Y, F379A)

· Mutated 5 (R194, D374Y, F379G)

· Mutated 6 (R194A, D374, F379)

· Mutated 7 (R194A, D374Y, F379)

· Mutated 8 (R194, D374, F379W)

· Mutated 9 (R194, D374, F379A)

The interactions were analyzed in terms of binding energy (kcal/mol) and chemical bonds.

Compounds and Reagents

The following compounds were selected for docking studies based on their potential to inhibit PCSK9: hypericin, amentoflavone, gingetin, bilobetin, and progesterone. These compounds were obtained from reputable suppliers and prepared for in silico analysis. The chemical structures of the compounds were verified using PubChem and ChemSpider to ensure accuracy in docking simulations.

Target Protein and Mutations

The structure of human PCSK9 (PDB ID: 3BPS chain A) was retrieved from the Protein Data Bank. The wild-type structure was utilized as the basis for subsequent mutagenesis studies. Key residues implicated in the binding to the LDLR EGF-A domain, namely Arg-194, Asp-374, and Phe-379, were targeted for mutation. The following mutations were introduced to generate the various PCSK9 variants using Chimera’s Rotamer Function (Shandler, 2010) [14]:

· Wild Type (R194, D374, F379)

· Mutated 1 (R194, D374Y, F379)

· Mutated 2 (R194A, D374Y, F379W)

· Mutated 3 (R194, D374Y, F379W)

· Mutated 4 (R194, D374Y, F379A)

· Mutated 5 (R194, D374Y, F379G)

· Mutated 6 (R194A, D374, F379)

· Mutated 7 (R194A, D374Y, F379)

· Mutated 8 (R194, D374, F379W)

· Mutated 9 (R194, D374, F379A)

Mutagenesis was performed using Molecular Docking Studies thorough Autodock Vina with Pyrx program [10,11].

Preparation of Protein Structures

The wild-type and mutant structures of PCSK9 were prepared for docking using the Pyrx program [10]. Water molecules and heteroatoms were removed, and hydrogen atoms were added to optimize the structure for docking simulations[10,11].

### 3.2 Preparation of Ligands

The chemical structures of the ligands were obtained in 3D format using Pubchem Server. The ligands were then minimized using MMFF94 Force Field (MMFF) to achieve a stable conformation suitable for docking.

#### Docking Procedure

Molecular docking simulations were performed using AutoDock Vina [10,11]. The following steps were taken:

Grid Box Definition: A grid box encompassing the binding site of PCSK9 was defined between PCSK9 and LDLR positioned at the coordinates of the critical binding residues (Arg-194, Asp-374, Phe-379) of PCSK9:

**a)** Wilde type

center_x = 24.0462651879; center_y = -32.5744876465, center_z = 1.51363427392;

size_x = 19.0057764664; size_y = 19.7149609362; size_z = 19.0057764664

**b)** Mutated 1 (D374Y)

center_x = 24.2265913404, center_y = -31.1009283831, center_z = 1.67330597378,

size_x = 17.7793572025, size_y = 15.8426372531, size_z = 16.5472745993

**c)** Mutated 2 (R194A, D374Y, F379)

center_x = 23.8218773228, center_y = -29.2913600211, center_z = 2.83341763807, size_x = 14.8956510965, size_y = 14.8956510965, size_z = 14.8956510965

d) Mutated 3 (R194, D374Y, F379W)

center_x = 23.3244242053, center_y = -30.8577951691, center_z = 0.75644361081, size_x = 18.2329935516, size_y = 16.6636036388, size_z = 18.2928598748

e) Mutated 4 (R194, D374Y, F379A)

center_x = 24.3432959999, center_y = -31.8004751101, center_z = 1.12608752883, size_x = 20.1995711195, size_y = 19.8898623855, size_z = 20.1995711195

f) Mutated 5 (R194, D374Y, F379G)

center_x = 23.4100970078, center_y = -31.3632022641, center_z = 2.01727491998, size_x = 16.3209069365, size_y = 16.3209069365, size_z = 16.3209069365

g) Mutated 6 (R194A, D374, F379)

center_x = 24.5878668913, center_y = -31.23572229, center_z = 1.30494752962, size_x = 19.0766267809, size_y = 14.9984321938, size_z = 17.3145523672

h) Mutated 7 (R194A, D374Y, F379)

center_x = 23.6946718945, center_y = -29.3349448548, center_z = 2.0355781923, size_x = 15.8312797284, size_y = 16.3564394597, size_z = 15.8312797284

i) Mutated 8 (R194, D374, F379W)

center_x = 23.1248602941, center_y = -31.1952343024, center_z = 0.308191091961, size_x = 19.1075573414, size_y = 18.9356693432, size_z = 19.5935839859

j) Mutated 9 (R194, D374, F379A) center_x = 23.1914553413, center_y = -30.0798973823, center_z = 0.0072884904863, size_x = 17.7165603888, size_y = 16.9031221894, size_z = 19.0533687587

Docking Protocol: The prepared ligand and receptor files were inputted into AutoDock Vina. The docking process was initiated, and 9 binding conformations were generated for each ligand-PCSK9 mutant complex.

Binding Energy Calculation: The binding energies of the resulting poses were calculated and ranked. The pose with the highest binding affinity (most negative binding energy) was selected for further analysis.

#### Molecular Interaction Analysis

The interactions between PCSK9 and the ligands were analyzed using the Discovery Studio software. Hydrogen bonds, hydrophobic interactions, and other non-covalent interactions were identified and characterized. The following interaction metrics were recorded for each docking pose:

· Number of Hydrogen Bonds: Counts of hydrogen bonds formed between the ligand and specific amino acid residues.

· Hydrophobic Interactions: Identification of hydrophobic contacts that contribute to the stability of the ligand-receptor complex.

· Key Interacting Residues: Notable residues involved in the binding interactions were highlighted for each ligand.

### Visualization and Reporting

All molecular structures, interactions, and binding modes were visualized using PyMOL [15] and Discovery Studio [16]. Detailed reports were generated to summarize the findings, highlighting key interactions and binding affinities for each ligand across the different PCSK9 mutants.

## Results

### Binding Affinity and Interactions

The docking results showed that Hypericin emerged as the most promising compound, maintaining significant binding affinity (kcal/mol) even in the presence of mutations. Notably, the analysis revealed that interactions between critical residues (Cys-378, Phe-379, and Arg-194) were essential for binding with Hypericin. The R194A mutation increased the affinity for some compounds while reducing it for others, suggesting that not all mutations lead to positive effects.

### Comparison Among Compounds

· Hypericin: Demonstrated the best binding affinity across almost all mutations, suggesting potential therapeutic utility in modulating PCSK9.

· Amentoflavone and Progesterone: Although they showed decent affinity, they did not reach the binding levels of hypericin.

· Bilobetin and Gingetin: Displayed variable binding affinity, with some mutations leading to unexpected results.

## Discussion

Docking analysis is a computational technique used to predict the interaction between a small molecule (ligand) and a target protein at the atomic level. This process involves finding the optimal position and orientation of the ligand within the binding site of the protein, based on binding affinity and interaction forces. The goal is to determine how well the ligand fits into the binding pocket and how strongly it binds [17–20].

In this study, we used docking analysis to evaluate the binding affinity of various natural compounds with both the wild-type and mutated variants of PCSK9. By doing so, we aimed to identify compounds that can effectively inhibit PCSK9-LDLR interactions, potentially preventing LDLR degradation and lowering LDL cholesterol levels (see table 1 below) . This approach allows us to explore potential therapeutic agents for hypercholesterolemia with minimal side effects[6]. The primary objective of this theoretical and complex study was to identify the key amino acids involved in the interaction between PCSK9 and the LDL receptor (as shown in Table 1). According to the study by Kwon et al. (2008), specific amino acids are responsible for this interaction, and mutating these residues enhances the binding affinity, leading to increased degradation of the LDL receptor (LDLR) and subsequent release of LDL into circulation [6].

**Table 1:**
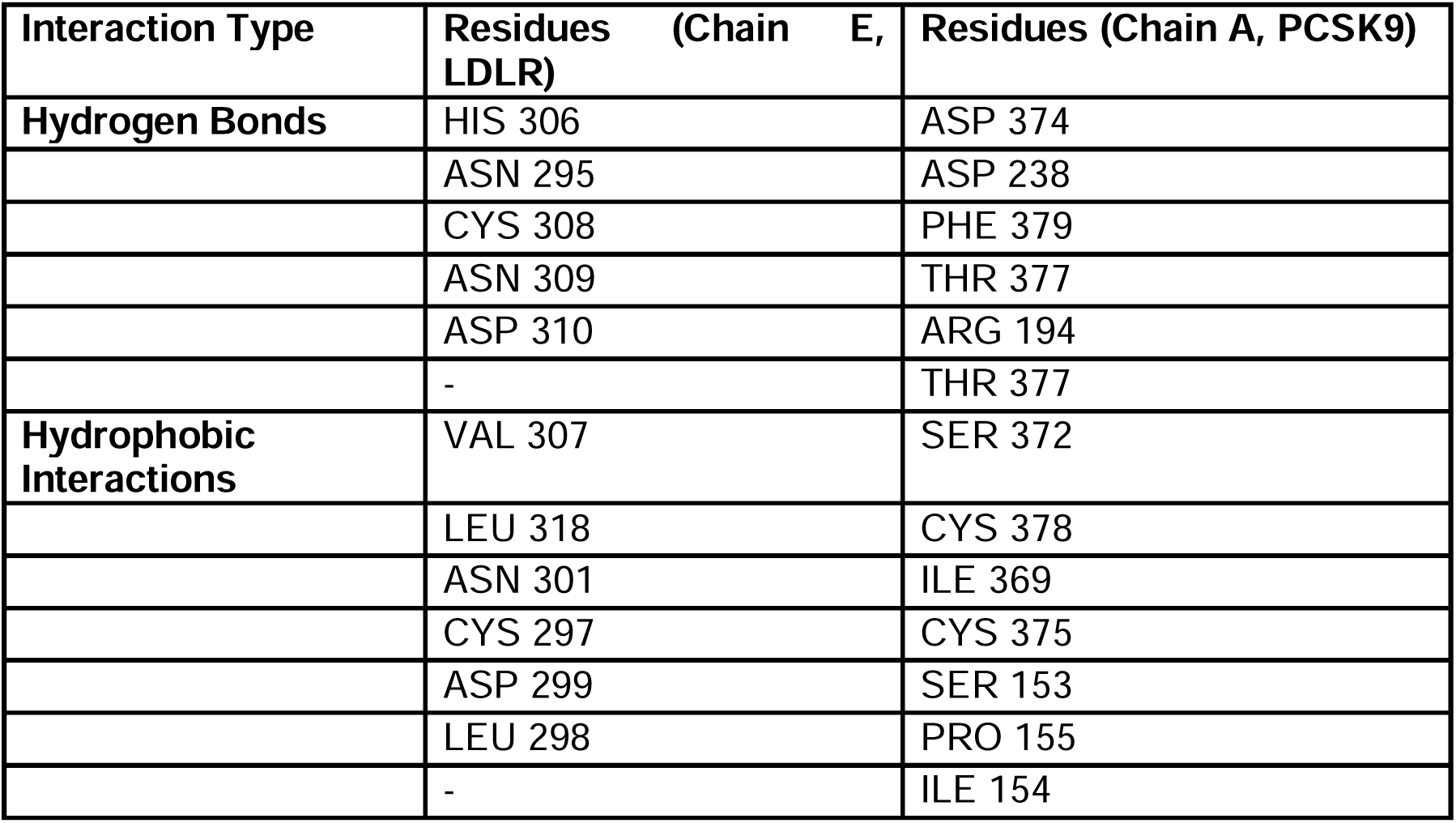
Comparison of Key Residues Involved Between LDLR and PCSK9 Calculated by LIGPLOT Program [21].

Following this understanding, a virtual screening was performed using the PyRx program [10]. A total of 100 natural compounds and some pharmaceuticals were docked into the active site of these key amino acids in PCSK9 using the AutoDock Vina software [11], and the binding energies (in kcal/mol) were calculated (see table 2 below).

**Table 2:** Comparison of Binding Energies (kcal/mol) of Natural Compounds and drugs in Selected Key Residue Zones of PCSK9 Proteins Calculated by Autodock Vina using Pyrx Program.

| Ligand | Binding Energy (kcal/mol) | Ligand | Binding Energy (kcal/mol) |
| --- | --- | --- | --- |
| 7-Dehydrocholesterol | -4.6 | Hesperetin | -5.2 |
| Alfacalcidol | -5.0 | Hesperidin | -5.8 |
| Apigenin | -5.2 | Hypericin | -7.3 |
| Artocarpin | -5.7 | Hyperoside | -5.3 |
| Ascorbic_Acid | -3.9 | Icariin | -5.0 |
| Aspirin | -4.2 | Icaritin | -5.9 |
| Astilbin | -4.9 | Isolicoflavonol | -5.3 |
| Astragalin | -5.5 | Isoorientin | -4.9 |
| Astringin | -5.5 | Isoquercetin | -5.4 |
| Baicalein | -5.4 | Isorhamnetin | -5.1 |
| Baicalin | -5.6 | Isovitexin | -4.9 |
| Bilobetin | -6.3 | Kaempferol | -5.0 |
| Blasticidin_S | -5.3 | Kanamycin | -4.2 |
| Caffeine | -3.8 | Kuromanin | -5.5 |
| Calcifediol | -5.2 | Luteolin_7-O-glucoside | -5.6 |
| Calycosin | -5.5 | Luteolin | -5.1 |
| Camphor | -3.1 | Marizomib | -4.1 |
| Capsaicin | -4.6 | Metronidazole | -4.3 |
| Cholecalciferol | -5.0 | Morin | -4.9 |
| Chrysin | -5.4 | Morusin | -6.1 |
| Coniferin | -4.4 | Mulberrin | -5.1 |
| Curcumin | -5.6 | Myricetin | -4.6 |
| Cyanidin | -5.0 | Myricitrin | -4.6 |
| Cyclomorusin | -5.8 | Naringin | -5.8 |
| Daidzein | -5.3 | Osajin | -5.8 |
| Daidzin | -5.5 | Papaverine | -5.1 |
| Diazepam | -5.9 | Paricalcitol | -5.4 |
| Diosmetin | -5.2 | Piceatannol | -5.5 |
| Diosmin | -5.5 | Pinosylvin | -5.4 |
| Disulfiram | -2.8 | Polydatin | -5.7 |
| Eleutheroside_B | -4.2 | Progesterone | -5.9 |
| Emodin | -5.2 | Quercetagetin | -5.1 |
| Equilin | -6.2 | Quercetin | -5.0 |
| Ergocalciferol | -6.1 | Quercitrin | -4.7 |
| Eriodictyol | -5.1 | Resistomycin | -6.2 |
| Estriol | -5.4 | Resveratrol | -5.7 |
| Estrone | -5.6 | Retinol | -4.8 |
| Fisetin | -5.2 | Rhapontin | -5.6 |
| Folic_Acid | -5.5 | Robustaflavone | -5.9 |
| Galangin | -5.3 | Romidepsin | -5.1 |
| Genistein | -5.1 | Rutin | -5.1 |
| Genistin | -5.3 | Sakakin_ | -5.2 |
| Ginkgetin | -6.3 | Sciadopitysin | -6.3 |
| Glabrone | -5.2 | Scutellarin | -5.5 |
| Gossypin | -5.2 | Shogaol | -4.2 |
| Spiraeoside | -4.8 | Silibinin | -5.5 |
| Squalene | -3.9 | Silymarin | -5.2 |
| Taxifolin | -4.9 | Ursolic_Acid | -5.0 |
| Tobramycin | -4.1 | Vanillin | -3.8 |
| Tricetin | -5.1 | Vitexin | -5.2 |
| Trifolin | -5.4 | amentoflavone | -6.5 |
|  |  | rottlerin | -5.7 |
|  |  | trans-Nerolidol | -4.3 |
|  |  | Bergamottin | -5.5 |

The five most promising compounds with the most negative binding energies, indicating a higher affinity for PCSK9, were selected for further analysis. These compounds included Hypericin, Amentoflavone, Bilobetin, Progesterone, and Gingetin.

Subsequently, these compounds were tested across nine different mutated PCSK9 structures, where the key amino acids were altered to evaluate their affinity and behavior in comparison to the wild type (R194, D374, F379). The goal was to determine if these compounds could act as potential inhibitors for PCSK9, which is known to cause LDL receptor degradation, leading to elevated circulating LDL levels. The rationale for mutating the key amino acids (R194, D374, F379) was to explore how alterations in these critical residues would affect the binding affinity of PCSK9 to the LDL receptor, as well as the efficacy of the natural compounds in inhibiting PCSK9 activity (see table 3) .

**Table 3:** Comparison of Binding Energies (kcal/mol) of Natural Compounds in Selected Key Residue Zones of PCSK9 Proteins calculated by Autodock Vina using Pyrx Program.

| Compound | Wild Type (R194, D374, F379) | Mutated 1 (R194, D374Y, F379) | Mutated 2 (R194A, D374Y, F379W) | Mutated 3 (R194, D374Y, F379W) | Mutated 4 (R194, D374Y, F379A) | Mutated 5 (R194, D374Y, F379G) | Mutated 6 (R194A, D374, F379) | Mutated 7 (R194A, D374Y, F379) | Mutated 8 (R194, D374, F379W) | Mutated 9 (R194, D374, F379A) |
| --- | --- | --- | --- | --- | --- | --- | --- | --- | --- | --- |
| Progesterone | -5,9 | -5,9 | -2,1 | -5,4 | -4,9 | -3,9 | -5,7 | -4,3 | -5,4 | -4,9 |
| Amentoflavone | -6,5 | -5,9 | 3,7 | -6 | -6 | -5,9 | -5,2 | -4,7 | -6,5 | -5,9 |
| Bilobetin | -6,3 | -4,5 | 8,1 | -5,9 | -6 | -5,1 | -6 | -4,1 | -6,1 | -6 |
| Gingetin | -6,3 | -5,7 | 3,3 | -5,2 | -5,9 | -5 | -6,1 | -3,5 | -5,9 | -5,8 |
| Hypericin | **-7,3** | **-6,4** | 19,8 | **-7** | **-5,9** | -4,9 | **-6,5** | -3,6 | **-7,2** | **-6** |

**Table 4:** Comparison of Hypericin with Mutated PCSK9 Structures (1-9) Compared to Wild Type (Reference)

| Wilde type of PCSK9 | Compounds | interactions |
| --- | --- | --- |
| R(194)D(374)F(379) | Hypericin ( -7.3 kcal/mol) | CYS (378),F(379),R(194) |
| mutated 1 |  |  |
| R(194)Y(374)F(379) | Hypericin ( -6.4 kcal/mol) | CYS (378),F(379),THR(377),R(194) |
| mutated 2 |  |  |
| A(194)Y(374)W(379) | Hypericin ( 19.8 kcal/mol) | CYS (378),CYS (375) |
| mutated 3 |  |  |
| R(194)Y(374)W(379) | Hypericin ( -7.0 kcal/mol) | TRP (379),ILE (369),THR(377) |
| mutated 4 |  |  |
| R(194)Y(374)A(379) | Hypericin ( -5.9 kcal/mol) | CYS (378),ILE (369),ALA(379) |
| mutated 5 | Compounds ( -4.9 kcal/mol) |  |
| A(194)D(374)F(379) | Hypericin ( -5.9 | PHE(379),CYS (378),THR (377) |
|  | kcal/mol) |  |
| **mutated 6** | Hypericin ( -6.5 kcal/mol) | PHE(379),CYS (378),THR (377) |
| A(194)D(374)F(379) |  |  |
| **mutated 7** |  |  |
| A(194)Y(374)F(379) | Hypericin ( -3.5 kcal/mol) | CYS (378),CYS (375),TYR (374),VAL (380) |
| **mutated 8** |  |  |
| R(194)D(374)W(379) | Hypericin ( -7.2 kcal/mol) | TRP (379),ILE (369),THR(377), CYS (378) |
| **mutated 9** |  |  |
| R(194)D(374)A(379) | Hypericin ( -6.0 kcal/mol) | CYS (378),ALA (379)ILE ( 369) |

The findings of this study underscore the importance of specific amino acid residues in PCSK9’s structure and their role in the protein’s interaction with LDLR. Our analysis revealed that Hypericin consistently demonstrated the highest binding affinity across nearly all mutations studied. This suggests that Hypericin may effectively inhibit PCSK9’s interaction with LDLR, making it a potential candidate for further development as a therapeutic agent. The critical interactions observed between hypericin and the residues Cys-378, Phe-379, and Arg-194 highlight the importance of these specific amino acids in maintaining strong binding affinities.

Interestingly, while some mutations, such as R194A, appeared to enhance binding for certain compounds, they also had a deleterious effect on others. This duality emphasizes the complexity of PCSK9’s binding dynamics and suggests that the therapeutic effectiveness of inhibitors may be influenced by the genetic background of patients. The implications of these findings are significant; for individuals with gain-of-function mutations in PCSK9, targeting the protein with a compound like hypericin could mitigate the adverse effects of such mutations on LDL metabolism.

The variable binding affinities observed for compounds such as amentoflavone and progesterone also warrant further exploration. Although these compounds did not match hypericin in terms of binding affinity, they still demonstrated considerable potential. Their varying interactions with mutated forms of PCSK9 indicate that these compounds may still be effective in specific contexts or patient populations. Future studies could explore the molecular mechanisms underlying these interactions in greater detail, potentially leading to the identification of novel therapeutic strategies.

Moreover, the structural dynamics of PCSK9, particularly the distance between its catalytic site and the binding surface for LDLR, suggest that targeting the interaction site may provide a therapeutic advantage without interfering with the catalytic activity of PCSK9. The development of small-molecule inhibitors that specifically disrupt the PCSK9-LDLR interaction, while preserving the protein’s physiological functions, could lead to significant advances in the treatment of hypercholesterolemia and its associated complication.

This docking analysis demonstrated that amino acid substitutions in PCSK9 significantly impact interactions with the studied compounds. Hypericin proved to be the most effective compound in maintaining binding affinity, indicating that it could be a promising candidate for further preclinical studies. Moreover, mutations in key residues, particularly R194 and D374, play a crucial role in the interaction between PCSK9 and LDLR, with potential implications for inhibitor development.

The key reason behind these mutations is to assess how reducing the size, charge, or hydrophobicity of the residues impacts the binding affinity of PCSK9 for LDLR. By disrupting these critical interactions, the aim is to reduce the overall strength of the PCSK9-LDLR complex. This allows for an understanding of which residues are most essential for maintaining this interaction and how small-molecule inhibitors like Hypericin might interfere with PCSK9’s function.

The rationale behind the mutations of Arg-194 and Phe-379 in PCSK9, as well as their impact on the interaction with the LDL receptor (LDLR), can be summarized as follows:

### Rationale for the Mutations

Arg-194

Arginine (Arg) is a positively charged amino acid with a large side chain that plays a significant role in maintaining polar interactions. Its mutation is aimed at reducing its size and charge, which are essential for stabilizing the interaction between PCSK9 and LDLR.

Arg-194 → Ala (R194A): Alanine is small and non-polar, leading to a loss of the positive charge and reduced steric bulk. This mutation is expected to weaken the interaction by eliminating key hydrogen bonds or electrostatic interactions that Arg- 194 forms.

Arg-194 → Ser (R194S): Serine is polar but uncharged. This substitution retains some potential for hydrogen bonding but reduces the strength of the interaction due to the absence of the positive charge.

Phe-379

Phenylalanine (Phe) is a hydrophobic, aromatic amino acid. It engages in π-π interactions and hydrophobic contacts, which stabilize the PCSK9-LDLR complex. Modifying this residue is expected to alter these critical interactions.

Phe-379 → Tyr (F379Y): Tyrosine retains the aromatic nature but introduces a hydroxyl group (-OH), which could form additional hydrogen bonds, potentially maintaining some interaction with LDLR.

Phe-379 → Trp (F379W): Tryptophan, with a larger aromatic ring system, might strengthen hydrophobic and π-π interactions, making it a strategic mutation to explore altered but not necessarily weaker binding.

Phe-379 → Ala (F379A): Replacing phenylalanine with alanine removes the aromatic ring entirely, reducing hydrophobic interactions and drastically weakening the binding affinity.

Phe-379 → Gly (F379G): Glycine is the smallest amino acid, removing steric bulk and completely eliminating aromatic interactions, likely reducing the stability of the complex.

Asp-374

Aspartate (Asp) is negatively charged and is known to be crucial for binding to LDLR. The mutation of Asp-374 to tyrosine (D374Y) is well-documented to increase PCSK9’s affinity for LDLR, leading to greater LDL receptor degradation. This mutation serves as a control to confirm how alterations in the binding site might affect compound efficacy.

These mutations serve two primary purposes:

Arg-194 and Phe-379 Mutations: To reduce the affinity between PCSK9 and LDLR by weakening or eliminating critical interactions.

Asp-374Y Mutation: To increase affinity, allowing a comparison between wild-type and enhanced binding states, helping identify whether certain compounds can still effectively inhibit PCSK9 under these conditions. Through these mutations, the goal is to find potential inhibitors that can reduce the degradation of LDL receptors, which in turn could lower circulating LDL levels.

From the analysis presented in Table 3, it is evident that Hypericin exhibits the highest binding affinity among the selected compounds, showcasing its superiority in interacting with PCSK9. This notable performance can be attributed to several factors:

Higher Binding Energy: Hypericin consistently displays more favorable binding energies compared to other compounds studied, indicating a stronger interaction with the PCSK9-LDLR complex. This suggests that Hypericin could effectively inhibit PCSK9’s function, which is crucial for regulating LDL cholesterol levels.

This table provides an insightful comparison, highlighting which residues are critical for the binding of selected natural compounds to PCSK9, offering potential leads for therapeutic inhibition.

Table 5 outlines the binding interactions of five selected compounds with different mutated structures of PCSK9 (See below) . The mutations focus on key amino acids (Arg-194, Asp-374, Phe-379), which play critical roles in the binding affinity between PCSK9 and LDLR. The reference wild-type (WT) structure is used for comparison to see how mutations alter binding affinity, and which amino acids are involved in the interaction for each compound. in the wild-type (WT) structure of PCSK9, the key residues Arg-194 (R194), Asp-374 (D374), and Phe-379 (F379) are critical for its interaction with the LDL receptor (LDLR). Here’s how each residue contributes:

- Arg-194 (R194): This residue plays a vital role in forming polar and electrostatic interactions. The positively charged side chain of arginine can establish hydrogen bonds and electrostatic interactions with negatively charged or polar residues of LDLR.
- Asp-374 (D374): This negatively charged residue is crucial for maintaining the binding affinity of PCSK9 to LDLR. The mutation of D374 to tyrosine (D374Y) has been shown to increase the affinity for LDLR in natural systems, leading to enhanced LDLR degradation.
- Phe-379 (F379): This aromatic, hydrophobic residue provides stabilizing interactions through hydrophobic contacts. It plays a role in maintaining the structural integrity and proper orientation of PCSK9 for binding to LDLR. In Mutated 1 (D374Y) of PCSK9, the critical mutation involves replacing Asp-374

**Table 5:**
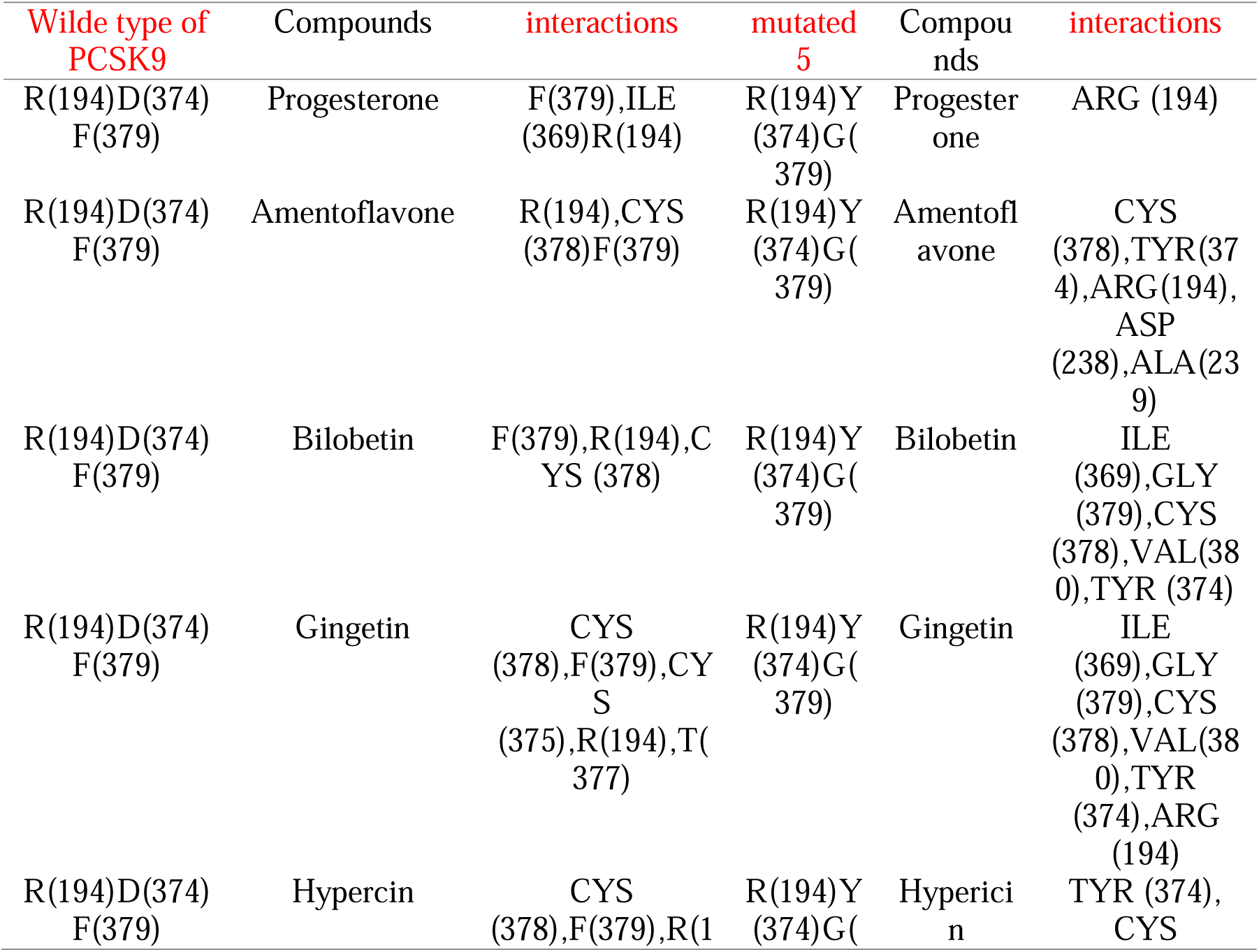
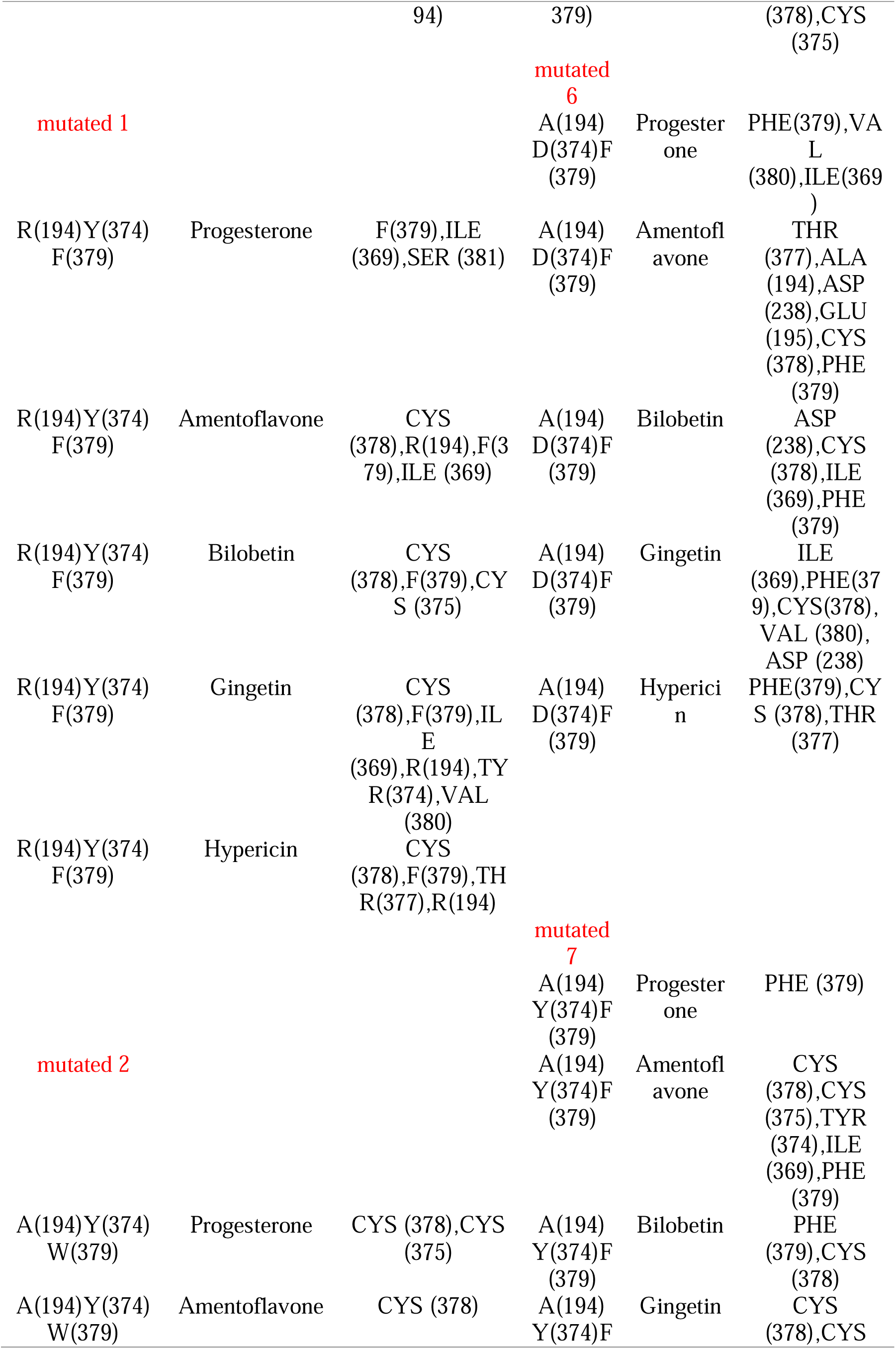
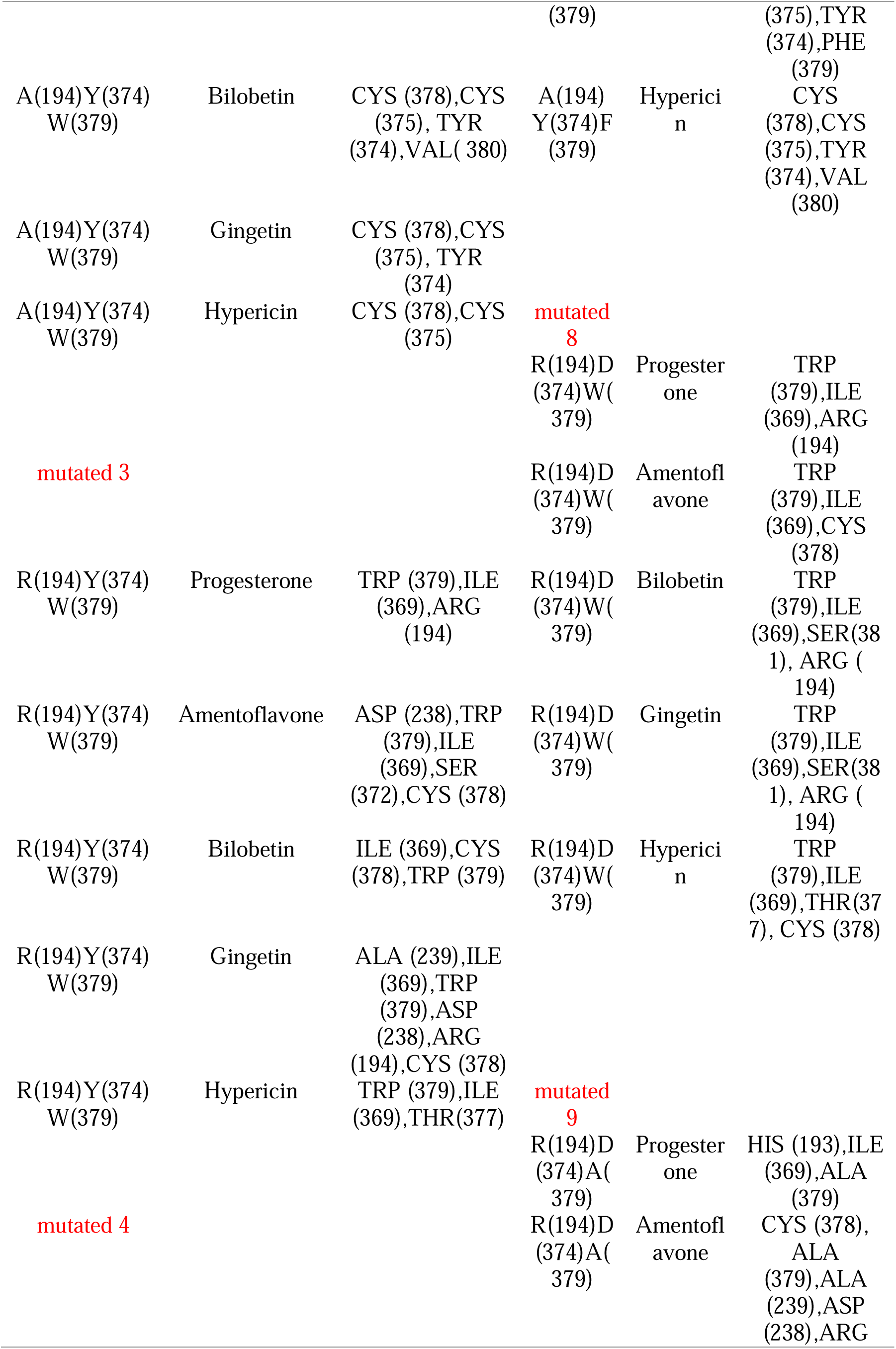
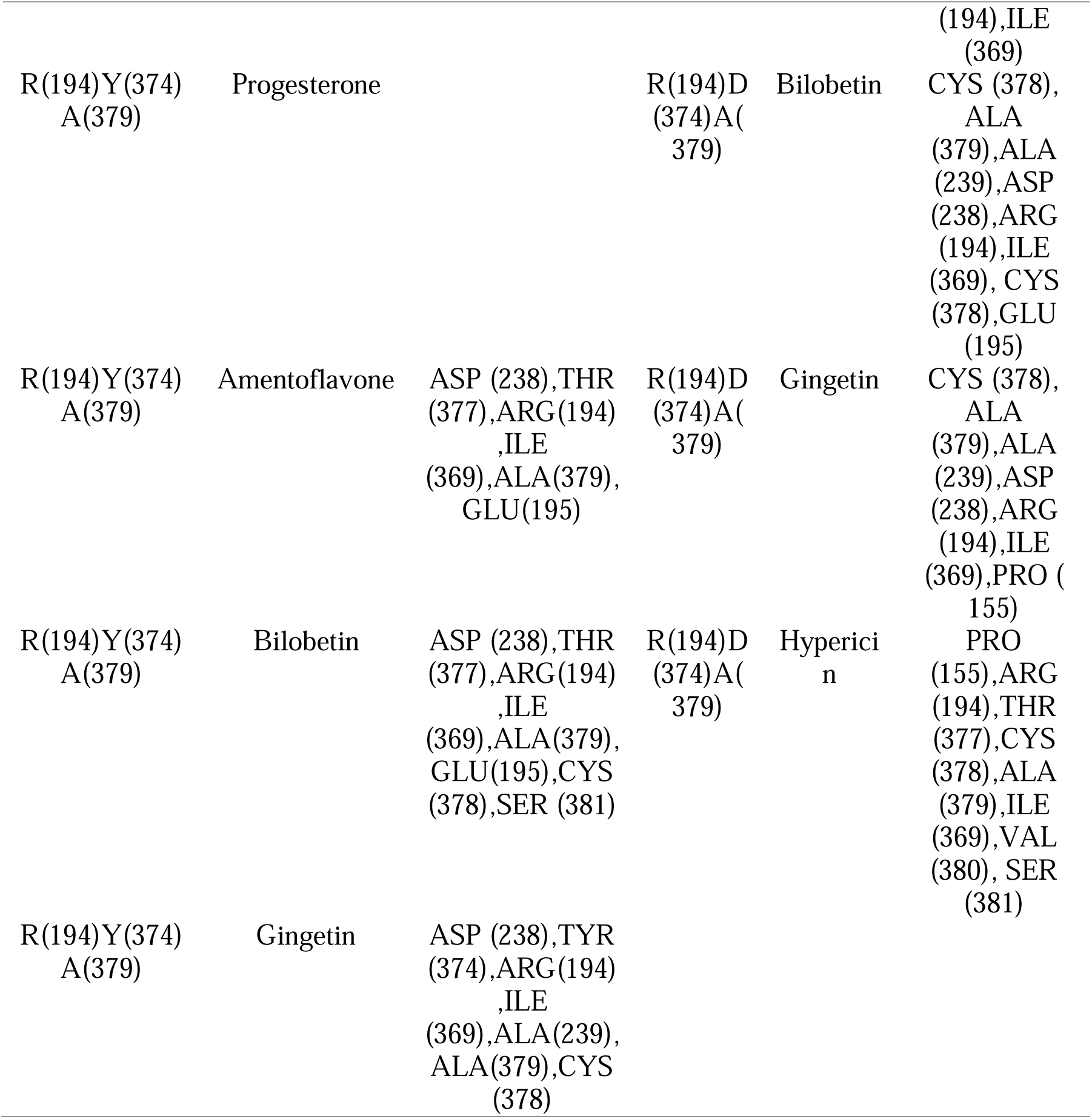
Comparison of selected compounds with Mutated PCSK9 Structures (1-9) Compared to Wild Type (Reference)

(D374) with tyrosine (Y), while Arg-194 (R194) and Phe-379 (F379) remain unchanged. Here’s how this mutation affects the structure and function:

- D374Y Mutation: The substitution of aspartic acid (Asp), which is negatively charged, with tyrosine (Tyr), a polar and aromatic residue, alters the local interaction dynamics. This mutation has been well-documented to increase the affinity of PCSK9 for the LDL receptor (LDLR). The reason for this increase is that tyrosine can participate in additional π-π stacking interactions or hydrophobic contacts that stabilize the PCSK9-LDLR complex. This leads to an enhanced degradation of LDLR, contributing to elevated LDL levels in circulation.
- Arg-194 (R194) and Phe-379 (F379): These residues retain their functions from the wild-type structure. Arg-194 continues to mediate polar and electrostatic interactions, while Phe-379 maintains its role in providing hydrophobic stabilization.

## Summary of Mutations and Implications

### Mutated 2 (R194A, D374Y, F379)

Mutations:

Arg-194 (R194) → Ala (A): Loss of positive charge reduces electrostatic interactions. Asp-374 (D374) → Tyr (Y): Increases affinity with LDLR.

Phe-379 (F379): Remains unchanged.

Implications: The overall binding affinity may slightly decrease due to the loss of Arg- 194’s charge, despite D374Y’s enhancement.

### Mutated 3 (R194, D374Y, F379W)

Mutations:

Asp-374 (D374) → Tyr (Y): Increases binding affinity.

Phe-379 (F379) → Trp (W): Maintains aromatic interactions, introducing bulk.

Implications: Increased steric bulk from Trp may alter binding dynamics, potentially maintaining strong interactions.

### Mutated 4 (R194, D374Y, F379A)

Mutations:

Asp-374 (D374) → Tyr (Y): Enhances binding affinity.

Phe-379 (F379) → Ala (A): Reduces hydrophobic interactions.

Implications: The benefits from D374Y may be balanced by reduced interactions from F379A, likely maintaining similar affinity to wild type.

### Mutated 5 (R194, D374Y, F379G)

Mutations:

Asp-374 (D374) → Tyr (Y): Enhances binding.

Phe-379 (F379) → Gly (G): Eliminates aromatic character.

Implications: The introduction of Gly could significantly weaken binding due to loss of stabilizing interactions.

*Mutated 6 (R194A, D374, F379*)

Mutations:

Arg-194 (R194) → Ala (A): Reduces positive charge. Asp-374 (D374): Remains unchanged.

Phe-379 (F379): Remains unchanged.

Implications: Reduced binding affinity from R194A may be offset by the stability of the unchanged residues.

### Mutated 7 (R194A, D374Y, F379)

Mutations:

Arg-194 (R194) → Ala (A): Reduces charge.

Asp-374 (D374) → Tyr (Y): Increases binding affinity. Phe-379 (F379): Remains unchanged.

Implications: D374Y may partially compensate for the loss of interaction from R194A, resulting in moderated binding affinity.

### Mutated 8 (R194, D374, F379W)

Mutations:

Asp-374 (D374): Remains unchanged.

Phe-379 (F379) → Trp (W): Increases interaction potential.

Implications: Retaining Asp-374 along with Trp may enhance overall binding through increased interaction possibilities.

### Mutated 9 (R194, D374, F379A)

Mutations:

Asp-374 (D374): Remains unchanged.

Phe-379 (F379) → Ala (A): Reduces aromatic interactions

Hypericin shows the highest binding affinity across both wild-type and most mutated structures (see table 3), particularly due to its ability to form strong interactions with key amino acids like Arg-194, Asp-374, and Phe-379. Mutations at Arg-194 generally lead to reduced affinity due to the loss of electrostatic and polar interactions. Phe- 379 mutations (particularly F379A and F379G) drastically decrease affinity as they remove the stabilizing hydrophobic interactions. The D374Y mutation, known to increase the affinity for LDLR in natural systems, results in only a slight reduction in binding affinity for the tested compounds, suggesting the compounds could remain effective even in the presence of this mutation., Hypericin demonstrates the highest binding affinity to PCSK9 across both wild-type and most mutated structures, largely due to its strong interactions with key residues such as Arg-194, Asp-374, and Phe- 379. Mutations at Arg-194 typically reduce binding affinity, as they disrupt important electrostatic and polar interactions. Similarly, mutations at Phe-379 (particularly F379A and F379G) cause a significant decrease in affinity by eliminating stabilizing hydrophobic interactions. Overall, Hypericin’s superior binding energy and its robust interaction with both wild-type and mutated amino acids of PCSK9 make it a promising candidate for further investigation as a potential therapeutic agent In Figures 1-3, the interactions of hypericin with involved residues in mutated PCSK9 (mutated 1-9) are shown and compared to the wild-type structure of PCSK9.

**Figure 1:**
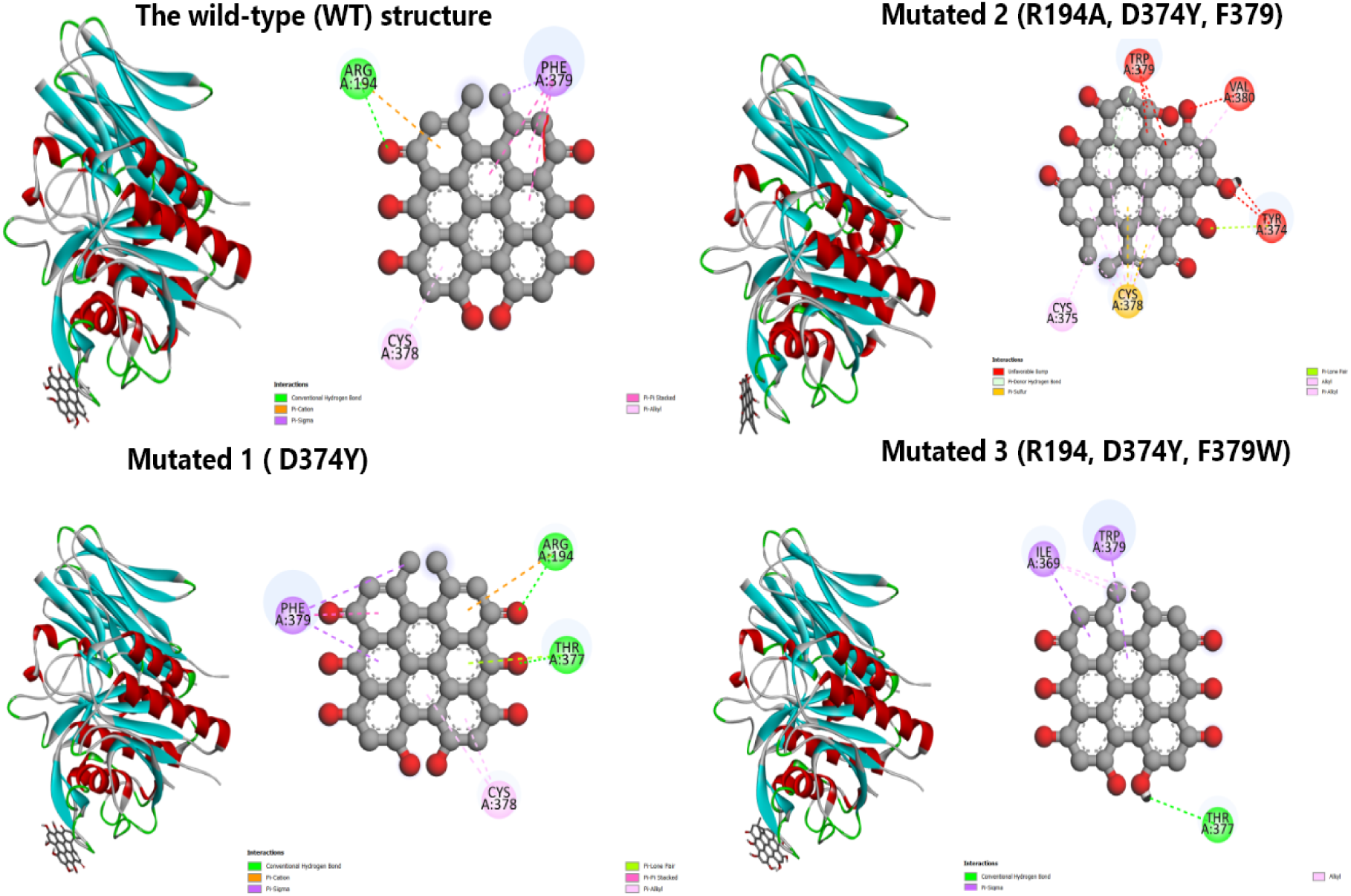
Comparison of Hypericin complexed with selected mutated regions of PCSK9 (from Mutated 1 to Mutated 3), using the wild type as a reference. Figure is performed by Discovery Studio Biovia Visualize

**Figure 2:**
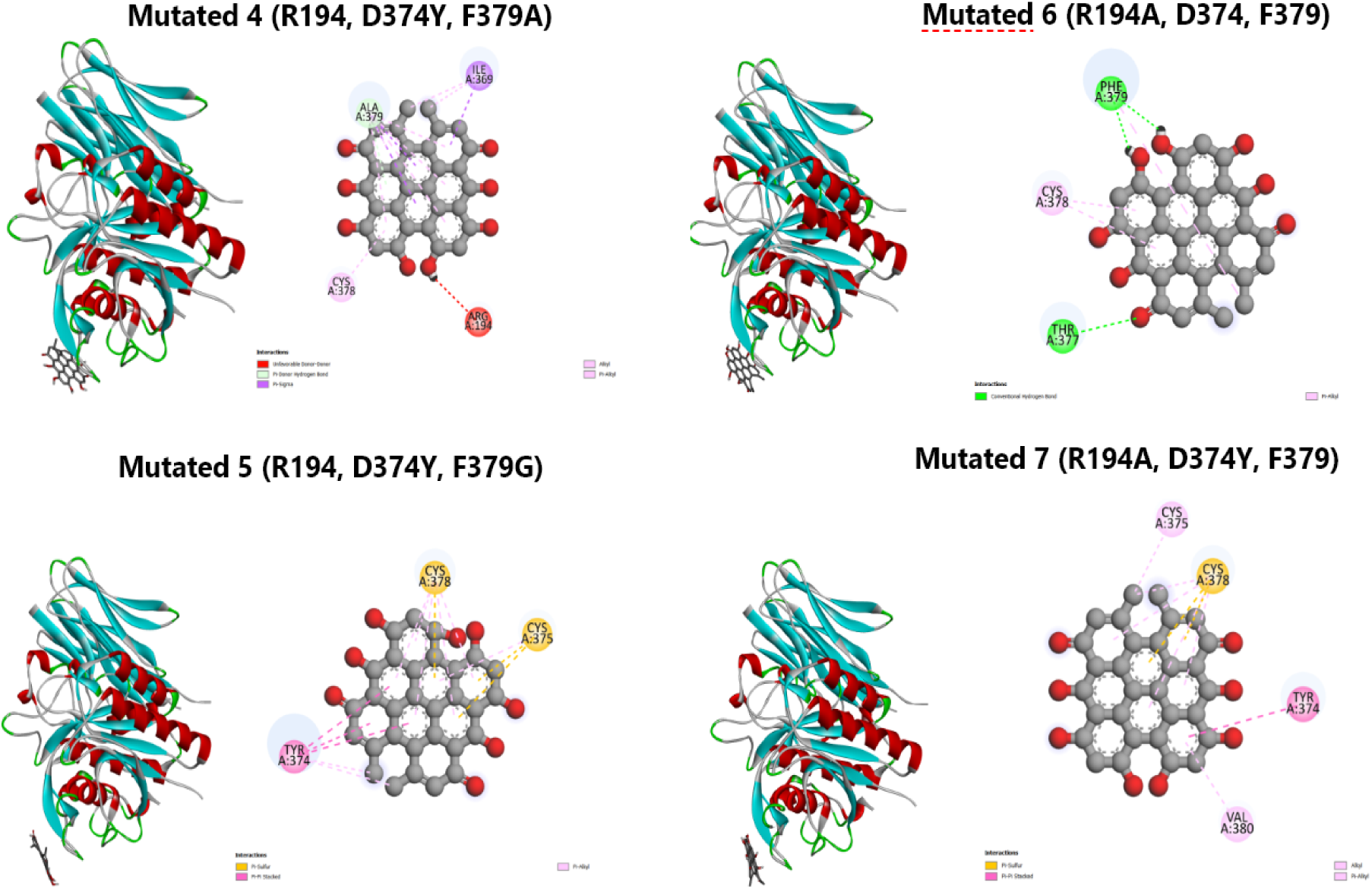
Comparison of Hypericin complexed with selected mutated regions of PCSK9 (from Mutated 4 to Mutated7), using the wild type as a reference. Figure is performed by Discovery Studio Biovia Visualized

**Figure 3:**
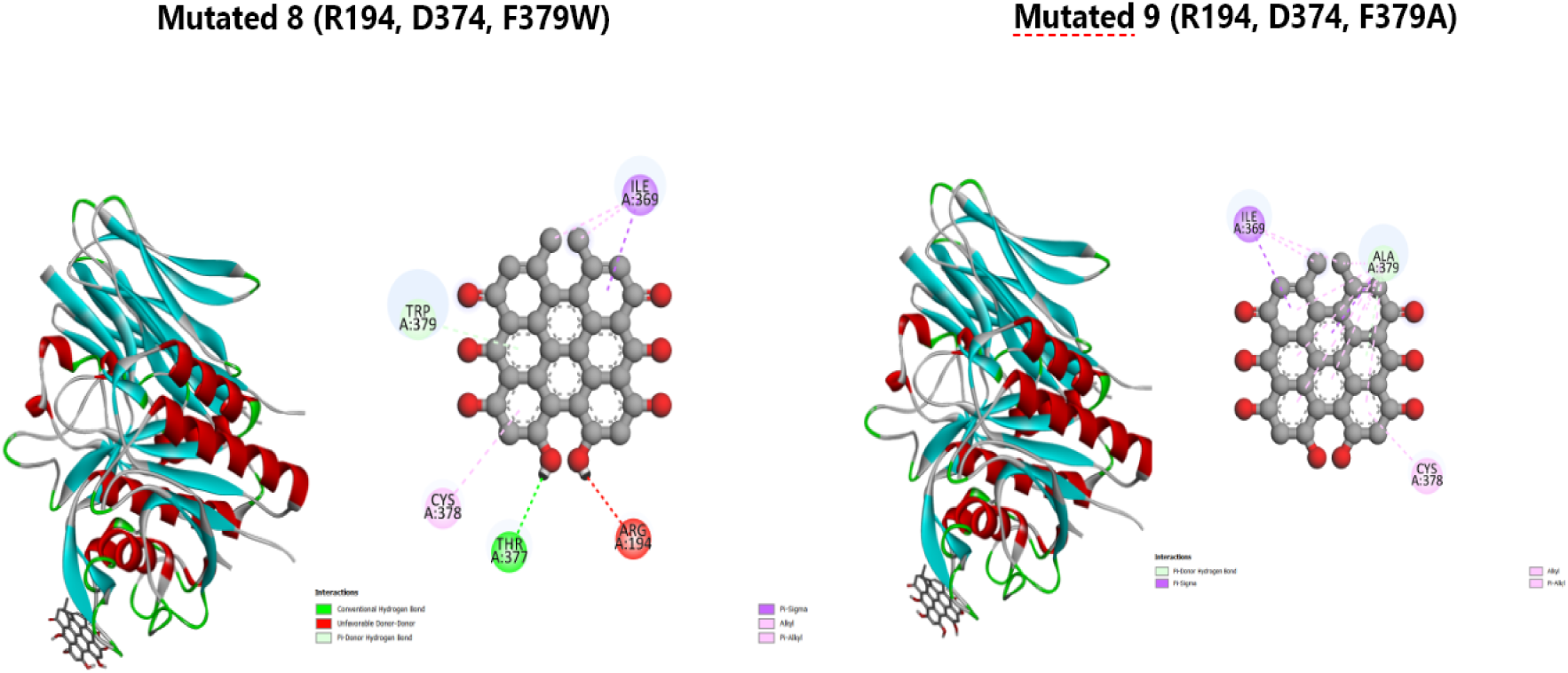
Comparison of Hypericin complexed with selected mutated regions of PCSK9 (from Mutated 7 to Mutated 9), using the wild type as a reference. Figure is performed by Discovery Studio Biovia Visualized

## Conclusion

Proprotein convertase subtilisin/kexin type 9 (PCSK9) plays a pivotal role in regulating cholesterol metabolism by promoting the degradation of low-density lipoprotein receptors (LDLR), leading to elevated LDL cholesterol levels in the bloodstream. Recent studies have identified key residues—Arg-194, Asp-374, and Phe-379—that are crucial for the interaction between PCSK9 and LDLR, and mutations in these residues can significantly alter this interaction. In this theoretical study, we employed molecular docking and in silico mutational analysis to assess the binding affinities of natural compounds, including Hypericin, Amentoflavone, Bilobetin, Gngetin, and Progesterone, to wild-type and mutant PCSK9. We explored nine specific PCSK9 mutations (R194, D374Y, F379), targeting changes in residues that influence LDLR binding affinity. Our in-silico approach reveals valuable insights into the effects of specific amino acid mutations on binding affinity, providing a foundation for the development of new therapeutic agents targeting PCSK9. As our understanding of the molecular mechanisms underlying cholesterol regulation deepens, we may unlock new strategies to combat hypercholesterolemia and reduce cardiovascular risk in affected individuals. Our results indicate that hypericin demonstrates favorable binding to wild-type PCSK9, interacting with the critical residues involved in LDLR binding, while other mutations yield diverse effects on compound affinities Hypericin demonstrated comparable binding energies in both the mutated and wild-type PCSK9 structures, suggesting its potential as a PCSK9 inhibitor. This compound could provide a novel approach to managing cholesterol levels and improving cardiovascular health, particularly in individuals with mutations that affect PCSK9 functionality. Despite these promising results, further theoretical studies, as well as in vitro and in vivo experiments, are needed to confirm its active biological role in blocking PCSK9 and preventing LDLR degradation.

The authors have declared no competing interest.

